# A CRISPR-based method for measuring the essentiality of a gene

**DOI:** 10.1101/736512

**Authors:** Yan You, Sharmila G. Ramachandra, Tian Jin

## Abstract

The CRISPR/Cas9 system is a powerful method of editing genes by randomly introducing errors into the target sites. Here, we describe a CRISPR-based test for gene essentiality (CRISPR-E test) that allows the identification of essential genes. Specifically, we use sgRNA-mediated CRISPR/Cas9 to target the open reading frame of a gene in the genome and analyze the in-frame (3n) and frameshift (3n+1 and 3n+2) mutations on the target region of the gene in surviving cells. If the gene is non-essential, the cells would carry both in-frame (3n) and frameshift (3n+1 and 3n+2) mutations. In contrast, the cells would carry only in-frame (3n) mutations if the targeted gene is essential, and this selective elimination of frameshift (3n+1 and 3n+2) mutations of the gene indicate its essentiality. As a proof of concept, we have used this CRISPR-E test in the model organism *Dictyostelium discoideum* to demonstrate that Dync1li1 is an essential gene while KIF1A and fAR1 are not. We further propose a simple method for quantifying the essentiality of a gene using the CRISPR-E test.

**One Sentence Summary:** CRISPR-E measures a gene’s essentiality

## Main Text

One of the most fundamental tasks of genetics is to identify genes that are essential for cellular or organismal viability ^1^. Current molecular genetic approaches, such as transposon mutagenesis, gene trapping, Restriction enzyme-mediated integration (REMI) mutagenesis, homologous recombination ^2^ and the more recently developed CRISPR gene editing ^3–8^, identify essential genes by first inactivating them and then determining if the cells or organisms carrying these mutations are not viable. Because these approaches use negative selection, there is no guarantee and often no proof that the gene has been disrupted or deleted (inactivated) in the dead cells. In the social amoeba *Dictyostelium discoideum*, a homologous-recombination method has been commonly used to knock out genes ^2^. However, we found that it is impossible to knock out some genes, and in those cases, one cannot conclude with certainty that these genes are essential due to the lack of evidence that the desired recombination events have occurred. To overcome this problem, we developed a simple CRISPR-based essentiality test to determine whether a gene is essential (named the CRISPR-E test) by targeting a gene of interest followed by analyzing mutations within the gene in surviving cells.

To evaluate essentiality of gene X using the CRISPER-E test (Fig. 1a), we express in cells the Cas9 protein and a sgRNA with a protospacer-adjacent motif (PAM) matching a location within gene X’s open-reading frame. Cas9 cuts DNA at the location to cause a double-strand break (DSB), and the DNA-repairing machinery fixes the break by non-homologous end joining (NHEJ), which ligates the ends and often introduces errors, including the insertion or deletion of a few nucleotides ^9–12^. If gene X is non-essential, surviving cells would carry alleles of WT and other mutations, which include both frameshift mutations (3n+1 and 3n+2, n=…−3, −2, −1, 0, 1, 2, 3…) that inactivate gene X and in-frame mutations (3n) that possibly retain the function of gene X. However, if gene X is essential, the cells would carry only in-frame mutations (3n) but no frameshift mutations (3n+1 and 3n+2). The presence of in-frame mutations proves that CRISPER/Cas9 has cut gene X and the DNA-repairing machinery has fixed the ends by non-homologous end joining, and thus, the selective elimination of frameshift mutations from living cells, a natural selection process, should indicate the essentiality of the gene.

**Fig. 1.**
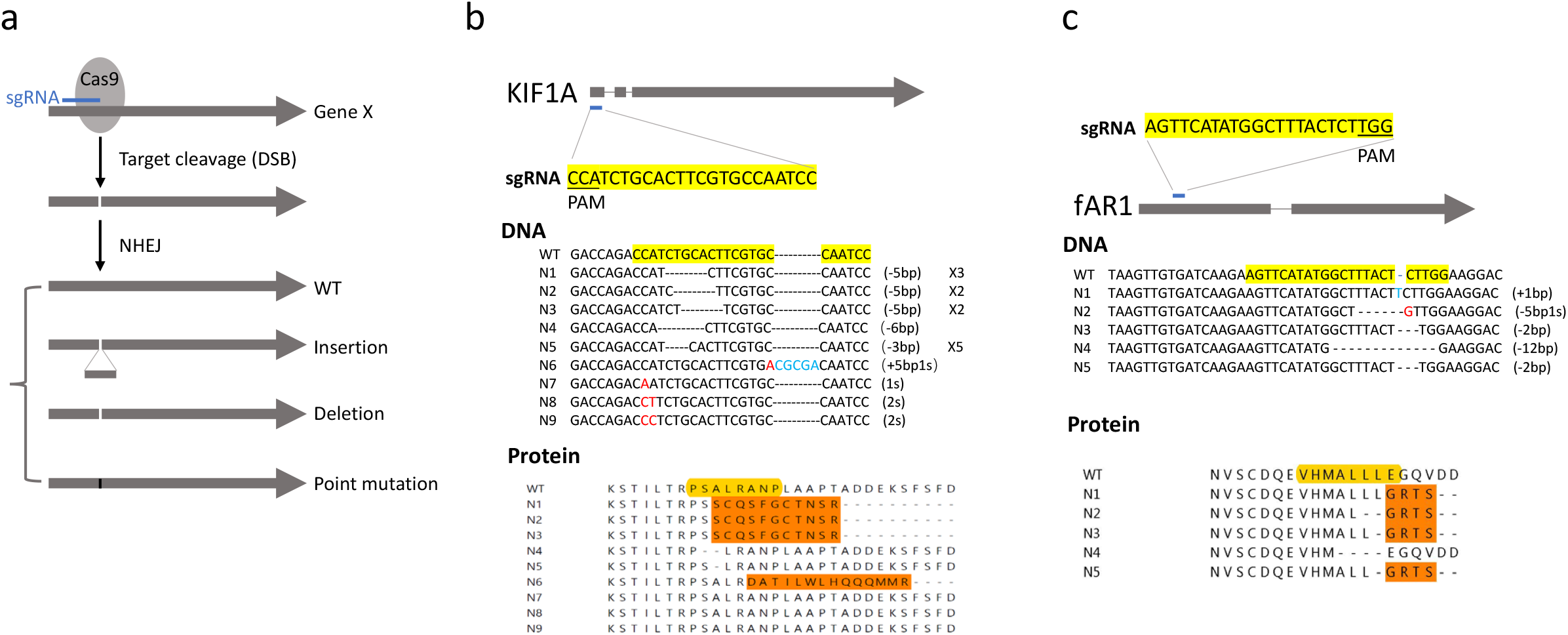
CRISPR-mediated gene editing. a. Mutational profiles of gene X generated by CRISPR/Cas9. A sgRNA matches a location and guides Cas9 to achieve a target cleavage at the location to cause double-stand break (DSB), followed by non-homologous end joining (NHEJ). After this sgRNA/Cas9-mediated gene editing, the DNA sequence on gene X should include WT, deletion, insertion, and point mutations. WT alleles are considered as alleles that have not been modified by CRISPR/Cas9 due to the lack of proof of gene editing, and other mutations are regarded as the products of gene editing. Edited alleles include WT, insertion, deletion and point mutations. Among them, frameshift mutations are 3n+1, 3n+2, and in-frame mutations are 3n (n=…-3, −2, −1, 0, 1, 2, 3…). b. A schematic view of sgRNA targeting KIF1A. The yellow-shaded sequence of sgRNA targets on the open reading frame of KIF1A. Under the label “DNA”, we show the sequencing results of the target regions from 19 individual clones (2WT, 3N1, 2N2, 2N3, N4, 5N5, N6, and N7, N8, N9). Red letter indicates a substation of a nucleotide, a dash shows a deletion of a nucleotide, and blue letter shows an insertion of a nucleotide. Under eh label “Protein”, we show the translated protein sequences of WT and N1-N9. N1, N2, N3 and N6 are frameshift mutations. The yellow-shaded sequence indicates the target site of sgRNA. c. A schematic view of sgRNA targeting fAR1. The yellow-shaded sequence of sgRNA targets on the open reading frame of fAR1. Under the label “DNA”, we show the sequencing results of the target regions from five individual clones (N1-N5). Under eh label “Protein”, we show the translated protein sequences of WT and N1-N5. N1-N4 and N5 are frameshift mutations.

To experimentally determine the effectiveness of the CRISPER-E test, we examined three genes, KIF1A (encoding kinesin 3), fAR1 (encoding folic acid receptor) and Dync1li1 (encoding dynein light intermedium chain) in *D. discoideum*. KIF1A and fAR1 have been knocked-out by a homologous recombination method and thus are not essential for cell proliferation ^13,14^, while Dync1li1 has not been knocked-out and has been suggested to be essential for the proliferation of *D. discoideum* cells ^15^. A CRISPR/Cas9 system has been developed to edit genes in *D. discoideum* ^16^. Using this system, we transformed cells with a vector expressing Cas9 and designed sgRNAs targeting on KIF1A (Fig. 1b), fAR1 (Fig. 1c) and Dync1li1 (supplementary Fig. 1 and Fig. 2), respectively. After the transformation of a vector of sgRNA/Cas9, the cells were plated on the SM agar with a bacterial lawn where they would grow as individual clones. We then randomly picked independent clones, amplified the target regions by PCR from each of the clones and sequenced the PCR products. Among 19 sequenced clones in which the sgRNA target was on KIF1A, two were WT, seven contain a 5-bp deletion at different positions, one contains a 6-bp deletion, five contain a 3-bp deletion, and, one contains a 5-bp insertion and a 1-bp substitution, and three contain point-mutations (Fig. 1b, DNA). There were eight frameshift mutations (3N1, 2N2, 2N3 and 1N6 in Fig. 1b, Protein), which should inactivate KIF1A function. Thus, the efficiency of on-target cutting/editing by sgRNA/Cas9 on KIF1A is about 89.5% (17 out of 19) and the efficiency of gene inactivation (frameshift mutations) is about 42% (8 out of 19). Among 5 sequenced fAR1 clones (Fig. 1c), there were four frameshift mutations and one in-frame mutation, and thus, the efficiency of on-targeting cutting/editing on fAR1 is about 100% and the efficiency of gene inactivation (frameshift mutations) is 80% (4 out of 5).

The result from the Dync1li1 gene is drastically different (Supplementary Fig. 1 and Fig. 2), and our CRISPR-E test indicates that Dync1li1 is an essential gene for cell growth. Specifically, we sequenced eight clones of Dync1li1 targeted by sgRNA1 and ten clones targeted by sgRNA2 (Supplementary Fig. 1) and found either wild-type or alleles with in-frame mutations (sgRNA1: four WT, two point mutations, one 3-bp deletion, one 3-bp insertion and one 21 bp deletion; sgRNA2: three WT, two point mutations, one 6-bp deletion, one 9-bp deletion, and three 3-bp deletions). While on-targeting cutting/editing efficiency were of 50% and 70% respectively, the efficiency of gene inactivation was 0. Assuming that CRISPR-mediated gene editing randomly generates 3n, 3n+1, and 3n+2 mutations, the probability of four and seven in-frame (3n) mutations by sgRNA1 and sgRNA2 are P1=(1/3)^4^=0.005 and P2=(1/3)^7^=0.00018, respectively, indicating that frameshift (3n+1 and 3n+2) mutations have been eliminated by natural selection during cell proliferation. To further increase the efficiency of cutting/editing on Dync1li1, we co-transformed vectors of sgRNA1/Cas9 and sgRNA2/Cas9 into cells and sequenced 21 individual clones. There are three WT and eighteen mutations, and the gene had been heavily edited on the region targeted by sgRNA1 and sgRNA2 with an 85% on-target efficiency (Fig. 2 DNA). However, each mutation was 3n that encodes a Dync1li1 protein with deletions, insertions or (and) substitutions of a few amino acids in the target region (Fig. 2 Protein), and no frameshift mutations were detected (Fig. 2 DNA), and thus, the efficiency of gene inactivation is 0. We interpreted the result as that frameshift mutations were eliminated because inactivating Dync1li1 is lethal to the cell proliferation process. This CRISPER-E test protocol allows us to identify an essential gene and determine the efficiency of gene inactivation, which provides a quantitative measurement of the essentiality of a gene.

**Fig. 2.**
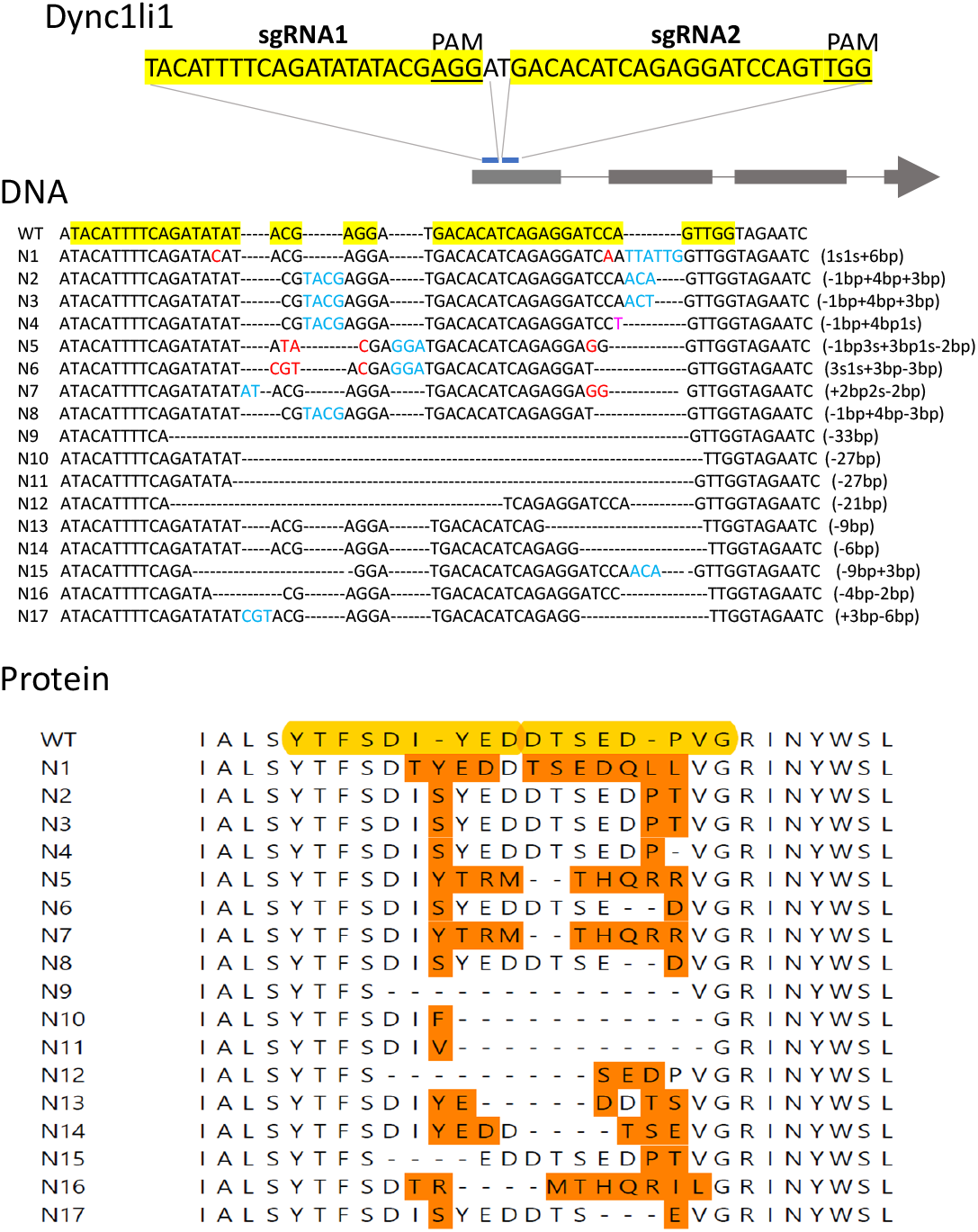
The CRISPR-E method shows that Dyncli1 is essential. A schematic view of sgRNA1 and sgRNA2 targeting Dync1li1 gene. The yellow-shaded sequences show sgRNA1 and sgRNA2 sequences targeting the open reading frame of Dyncli1. Under the label “DNA”, we show the sequencing results of the target regions of 21 individual clones, which includes 3WT, N1-14, 2N15, N16 and N17. Red letter indicates a substation of a nucleotide, a dash shows a deletion of a nucleotide, and blue letter shows an insertion of a nucleotide. Under the label “Protein”, we show translated protein sequences of WT and N1-N17. Yellow-shaded sequences show the regions targeted by sgRNA1 and sgRNA2. All mutant alleles are in-frame (3n) mutations.

We propose that the CRISPR-E test can quantitatively measure essentiality of all genes. CRISPR-based gene cutting/editing efficiency is high and can be measured experimentally. Cas9-geneated mutations from imperfect action of NHEJ randomly create frameshift (inactivating) mutations and in-frame mutations. The theoretical frequency of mutations on sgRNA/Cas9 on a target site of a gene can be calculated using mathematical models and computing programs, and these methods can be further developed in future. Experimentally measured frequency of gene inactivation (frameshift mutations of a gene) divided by the theoretical frequency gives rise to a number E, which provides a measurement of a gene’s essentiality. E>1 means that inactivating the gene is bifacial for cell’s survival, E=1 means that inactivating the gene has no effect on survival, 1>E>0 means that inactivating the gene has a negative effect on survival, and T=0 means that the gene is essential. CRISPR-E test is a simple method that can be applied to quantitatively measure the essentiality of every gene in haploid or diploid cells of all organisms. If a gene is not essential for cell growth and proliferation, its essentiality can be further examined in other biological process, such as cell migration, cell differentiation, and development of multicellular organisms. Furthermore, essentiality of genes can be measured and screened in the development of human diseases in cell culture or animal models. For example, it will be particularly useful to determine which genes are highly or completely essential (with E score very low or E=0) for the proliferation or survival of cancer cells but are less or not essential (with E score very high) for that of normal cells. The proteins encoded by those genes are likely to be good targets of cancer therapeutic drugs. Future research efforts are required to quantitatively measure the essentiality of genes of interest and to screen for those that are critical for various biological processes, by using this simple CRISPR-E method in different genetic background, environments and organisms.

## Methods

Strains, cell culture, plasmid construction and PCRs. *Dictyostelium discoideum* strains AX2 (wild type) and their transformants were incubated at 22°C on culture dishes or in shaking culture in HL5 medium or on SM agar with *Klebsiella aerogenes*. Transformants were selected in HL5 medium with G418 at 20 μg/ml. Using pTM1285 as the parental plasmid ^16^, we inserted a gRNA scaffold-sgRNA sequence into sgRNA expressing cassettes and generated vectors expressing Cas9, GFP, G418 resistance cassette and sgRNA targeting KIF1A, fAR1 or Dync1li1 genes. Each of the plasmids was transformed into AX2 (wild-type) cells. To sequence the target regions edited by sgRNA/Cas9 in individual clones, we amplified genomic DNA using PCR with the following primers: KIF1AF (5’-GAAGAGCAAGGTAAAAAGG-3’), KIF1AR (5’-CCTTTTACCAGAACCAGTTTG-3’), and the PCR product is 400bp for WT; for fAR1, fAR1F (5’-ACGACCCATTGTATTAT-3’), fAR1R (5’-GACTTTGAGTACA AATATCG-3’), and the PCR product is 530 bp for WT; for Dync1li1, Dync1li1(5’-AAGAAGATATTTGGGGTC-3 ‘), Dync1li1R (5’-GGTTGTGAAAAATCTAAAG-3’), and the PCR product is 337bp for WT.

## Acknowledgments

We thank Tetsuya Muramoto from Toho University, Chiba, Japan for providing the pTM1285 plasmid, Xuehua Xu and Miao Pan for helpful discussions, and Xin Xiang for a critical reading of the manuscript. The work is supported by intramural funding from NIAID, NIH.

---

**Supplementary Fig. 1.**
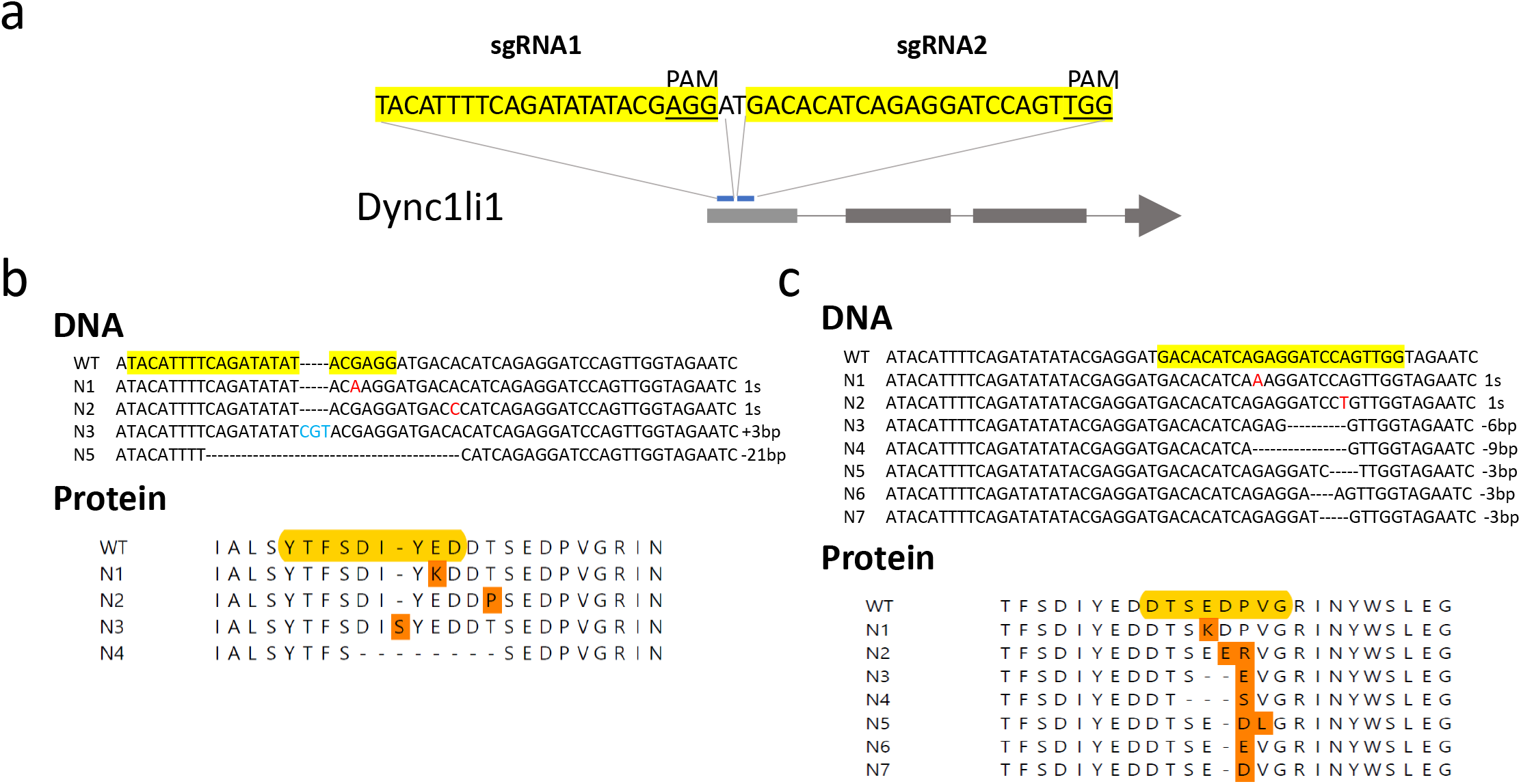
**a.** A schematic view of sgRNA1 and sgRNA2 targeting Dync1li1 gene. The yellow-shaded sequences show sgRNA1 or sgRNA2 sequences targeting the open reading frame of Dyncli1. **b.** Dync1li1 gene edited by sgRNA1. Under the label “DNA”, we show the sequencing results of the target regions of eight individual clones, which includes 4WT, N1-N4. Red letter indicates a substation of a nucleotide, a dash shows a deletion of a nucleotide, and blue letter shows an insertion of a nucleotide. Under the label “Protein”, we show the translated protein sequences of WT and N1-N4. Yellow-shaded sequences show the regions targeted by sgRNA1. c. Dync1li1 gene edited by sgRNA2. Under the label “DNA”, we show the sequencing results of the target regions of ten individual clones, which includes three WT, N1-N7. Red letter indicates a substation of a nucleotide, a dash shows a deletion of a nucleotide, and blue letter shows an insertion of a nucleotide. Under the label “Protein”, we show the translated protein sequences of WT and N1-N7. Yellow-shaded sequences show the regions targeted by sgRNA2. All mutant alleles are in-frame (3n) mutations.

